# Individual recognition in a jumping spider (*Phidippus regius*)

**DOI:** 10.1101/2023.11.17.567545

**Authors:** Christoph D. Dahl, Yaling Cheng

## Abstract

Individual recognition is conceptually complex and computationally intense, leading to the general assumption that this social knowledge is solely present in vertebrates with larger brains, while miniature-brained animals in differentiating societies eschew the evolutionary pressure for individual recognition by evolving computationally less demanding class-level recognition, such as kin, social rank, or mate recognition. Arguably, this social knowledge is restricted to species with a degree of sociality (sensu [1], for a review [2]). Here we show the exception to this rule in a non-social arthropod species, the jumping spider *Phidippus regius*. Using a habituation - dishabituation paradigm, we visually confronted pairs of spatially separated spiders with each other and measured the ‘interest’ of one spider towards the other. The spiders exhibited high interest upon initial encounter of an individual, reflected in mutual approach behaviour, but adapted towards that individual when it reoccurred in the subsequent trial, indicated by their preference of staying farther apart. In contrast, spiders exhibited a rebound from habituation, reflected in mutual approach behaviour, when a different individual occurred in the subsequent trial, indicating the ability to tell apart spiders’ identities. These results suggest that *P. regius* is capable of individual recognition based on long-term memory.

## Main text

Recognising individuals is a complex cognitive process requiring flexible learning and recognition memory. Arthropod species possessing the social ability of individual recognition would, thus, stand in stark contrast to the commonly accepted notion that animals with smaller brains are cognitively less advanced due to reduced computational power of nervous systems with smaller and fewer neurons [3]. And yet, there is evidence for an arthropod species displaying face learning [4] and long-term social memory [5]. That is, a social wasp species (*Polistes fuscatus*) showed mammal-like face learning [4, 6], arguably providing social benefits by reducing aggression and stabilizing social interactions. As one of the few reported instances of individual recognition in arthropods (see also [7]), this has contributed to the prevailing view that non-social arthropod species are unlikely to evolve such complex cognitive processes. The underlying reasoning is that individual recognition entails high energetic costs, longer processing times, and consequently an increased risk of predation – costs that would not be outweighed by the limited number of social encounters or the marginal survival benefits they provide [2, 8]. The general consensus, thus, is that a certain degree of sociality sensu Wilson [1] is required for the emergence of individual recognition [8]. Here, we challenge this consensus by testing for individual recognition in *Phidippus regius*, a non-social, miniature-brained jumping spider. In a controlled experimental procedure, we confronted subjects with live conspecifics, intensifying the social encounter and ensuring ecological relevance in the stimulus presentation. While jumping spiders have been shown to use visual and chemical cues for species, sex, or rival recognition [9, 10], there is no direct evidence for memory-based recognition of individual identity, particularly in taxa such as jumping spiders, where repeated encounters with the same conspecific are infrequent.

As a first step, we assessed the ability of *P. regius* to individually recognise other members of its species, commonly referred to as individual recognition [11] or individuation of conspecifics [12]. To test this, we employed a habituation - dishabituation paradigm, in which one individual habituates to the extended presence of another individual in its close proximity. Following a brief phase of visual separation, a different individual is introduced. If the focal individual can discriminate the current individual from the former, it is expected to exhibit dishabituation [13, 14]. In other words, in this habituation - dishabituation paradigm, we expect the rebound in ‘interest’ to be greater when the identity of the spider changes than when the same individual is presented again. To experimentally control the animal pairs, we placed each individual in a separate container with one transparent side and a transparent top panel. We then pairwise confronted the individuals by placing the containers such that the transparent sides faced each other, allowing the spiders to visually explore and approach one another at close range while preventing any direct physical interaction.

Each trial followed the same procedure: Two spiders, say individuals A and B, were exposed to one another for 7 minutes, eliciting an initial ‘interest’ in each other. They were then visually separated for 3 minutes using an opaque slider. Following this separation, the same pair could be re-exposed to one another (A vs B, *habituation* trial) or either individual could be paired with a new individual (A vs C, or B vs D, *dishabituation* trial), again for 7 minutes, followed by another 3-minute separation period. Relative interest was quantified by approximating spatial distances between the spider pairs in the xy-plane from video recordings captured through the transparent top panel, where high interest is reflected in smaller values (i.e. spiders move closer) and low interest in larger values (spiders maintain distance). Under the assumption that spiders are capable of individuating each other, we predict that in the *habituation* condition, where the same individuals are re-encountered, relative interest decreases, resulting in an increase in distance (Figure 1a-c; ‘Habituation’, hypothetical example 1: spiders seek maximal distance to each other (dashed line); example 2: spiders seek medium distance to each other (solid line)). Conversely, in the *dishabituation* condition, where a new individual is introduced, relative interest increases and spiders approach each other, leading to a decrease in distance (Figure 1a-c; ‘Dishabituation’). It is important to clarify the use of the term ‘Baseline’ as illustrated in Figure 1a. The *baseline* trial shown there represents only the first trial in each session, before any habituation or dishabituation has occurred. For all subsequent comparisons, relative changes in proximity were computed by comparing each trial to the immediately preceding one. Specifically, each *dishabituation* trial was analysed relative to the preceding *habituation* trial (Figure 1c: ‘Dishabituation - Habituation’), while each *habituation* trial (except for the first in each session) was analysed relative to the preceding *dishabituation* trial (Figure 1c: ‘Habituation - Baseline’, equivalent to ‘Habituation - Dishabituation’). Thus, habituation and dishabituation are not absolute responses, but are defined by how much the behaviour shifts relative to a reference point. As a result, the difference scores reflect relative changes in the frequency distribution of distances. Some values in the hypothetical distributions (Figure 1c) are negative, indicating that the number of times a spider was at a given distance was lower compared to the preceding trial. A decrease in interest can be inferred when this underrepresentation (i.e. negative values) occurs in the more proximal distance bins, accompanied by an overrepresentation (i.e. positive values) in the medium or distal distance bins. Conversely, an increase in interest can be inferred when there is an overrepresentation of occurrences (i.e. positive values) in the proximal distance bins, coupled with an underrepresentation (i.e. negative values) in the medium or distal distance bins. These relative changes in proximity describe the behavioural signature of habituation and dishabituation (Figure 1c) and constitute the basis for our statistical analyses.

**Figure caption 1:**
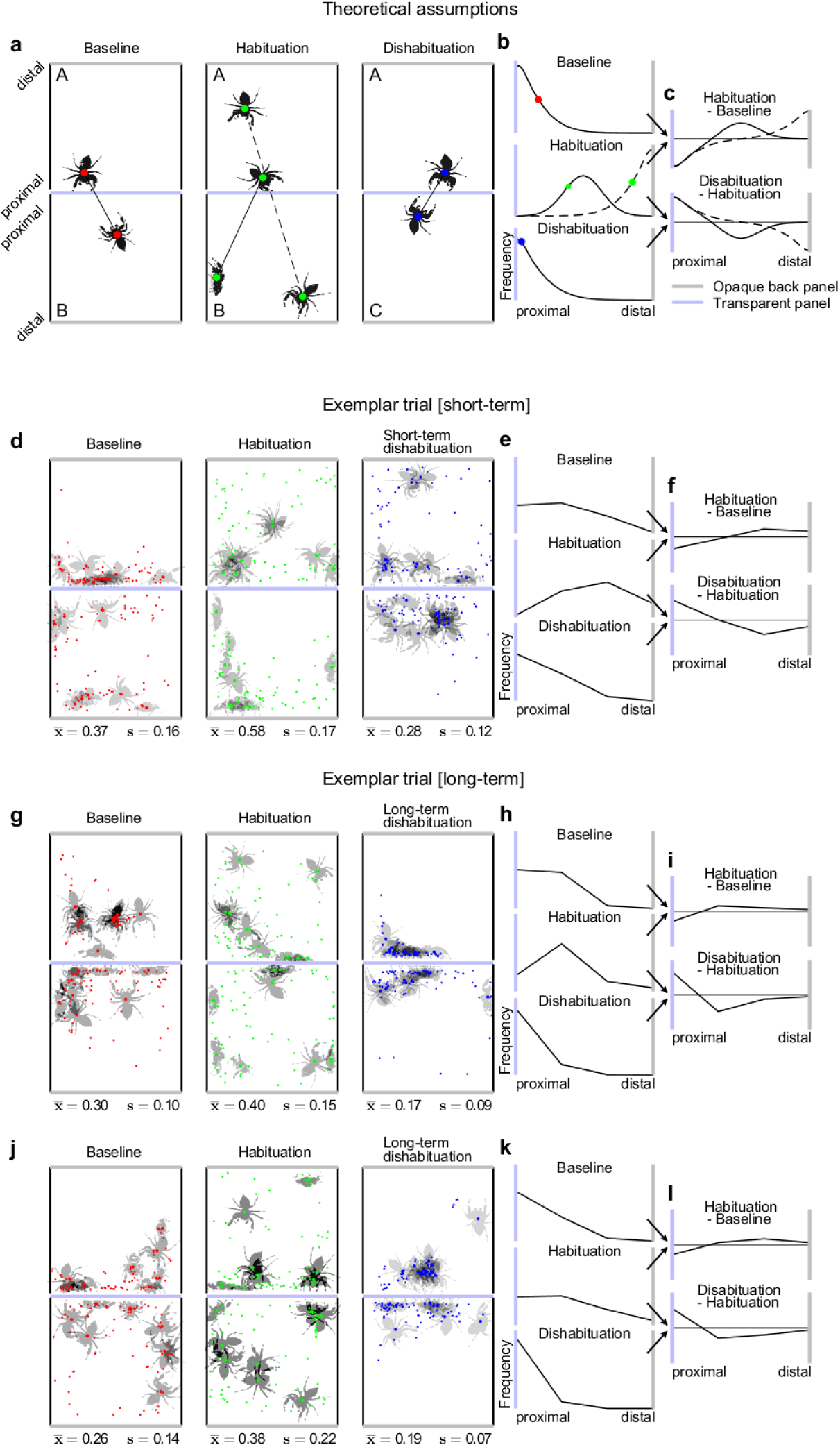
Theoretical assumptions and exemplar trials. a-c Predicted spider distances for *baseline* (red dots), *habituation* (green dots) and *dishabituation* comparisons (blue dots). Habituation can manifest either in equal inter-spider distances (solid line) as in the *baseline* comparison or in an increase of distances (dashed line). What is referred to as *baseline* in this context is the *dishabituation* trial of the previous comparison (see Table 2). Distance samples are predicted to fall into distributions as shown in b. Contrasts between *baseline*, *habituation* and *dishabituation* comparisons would result in distributions as shown in c. d-f An exemplar trial consisting of *baseline*, *habituation* and *dishabituation* comparisons from the first session of trials is shown. The *short-term dishabituation* comparison shows a decrease of inter-spider distances, indicating increasing interest in a different individual than the previously perceived one (*habituation* comparison). g-i; j-l Two exemplar trials from the third session of Experiment 2 are shown, where a presentation of an individual novel and unseen across the three experimental sessions triggered a great rebound in interest (i,l, ‘Dishabituation - habituation’). d, g, j: Note that in all upper quadrants, the same spider is used for baseline, habituation, and dishabituation comparisons, while in the lower quadrants, the baseline and habituation involve one individual, and a different (novel) spider is presented in the dishabituation comparison.

In the first experiment, we divided a total of 20 individuals into five groups of four individuals each. Each individual of each group was exposed to the three group members in both habituation and dishabituation trials, resulting in six trials per session, equivalent to one hour of recording. We repeated this procedure twice, resulting in 18 trials across three sessions and an exact repetition of a given trial (and pairing of individuals) in 1-hour intervals (for a detailed description of the procedure see Materials and Methods and Tables 1-2).

**Table 1:** Pairwise comparisons.

| Trial | Pair 1 | Pair 2 |
| --- | --- | --- |
| 1 | A - B | C - D |
| 2 | A - C | B - D |
| 3 | B - C | A - D |

**Table 2:**
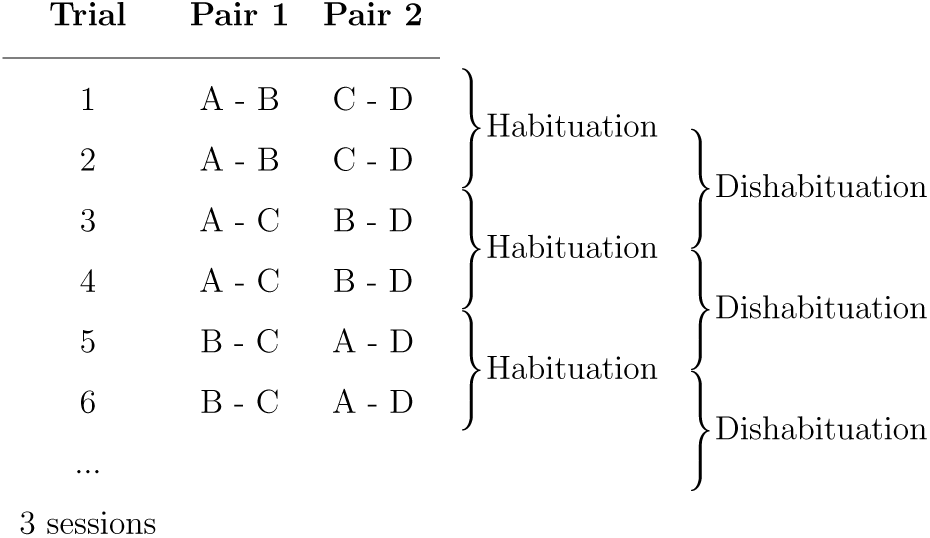
Procedure of Experiment 1.

We found that spiders adjusted their proximity depending on whether they encountered a familiar or a new individual. In *habituation* trials, where the same individual was presented again, spiders tended to maintain greater distances. In contrast, in *dishabituation* trials, where a new individual was introduced, spiders were more likely to stay at close distances. This pattern was statistically robust: *habituation* and *dishabituation* trials (i.e. predictor variable *condition*) differed significantly as a function of inter-individual distances (i.e. predictor variable *distance*), leading to a significant improvement of model fitting the interaction of the predictors *distance* and *condition* (LRT: 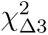 = 63.66, *p* < 0.001; Figure 2a, Supplementary Table 1): *dishabituation* trials (blue discs) showed a greater proportion of close-distance values than *habituation* trials (red discs), whereas *habituation* trials showed a greater proportion of far-distance values.

**Figure caption 2:**
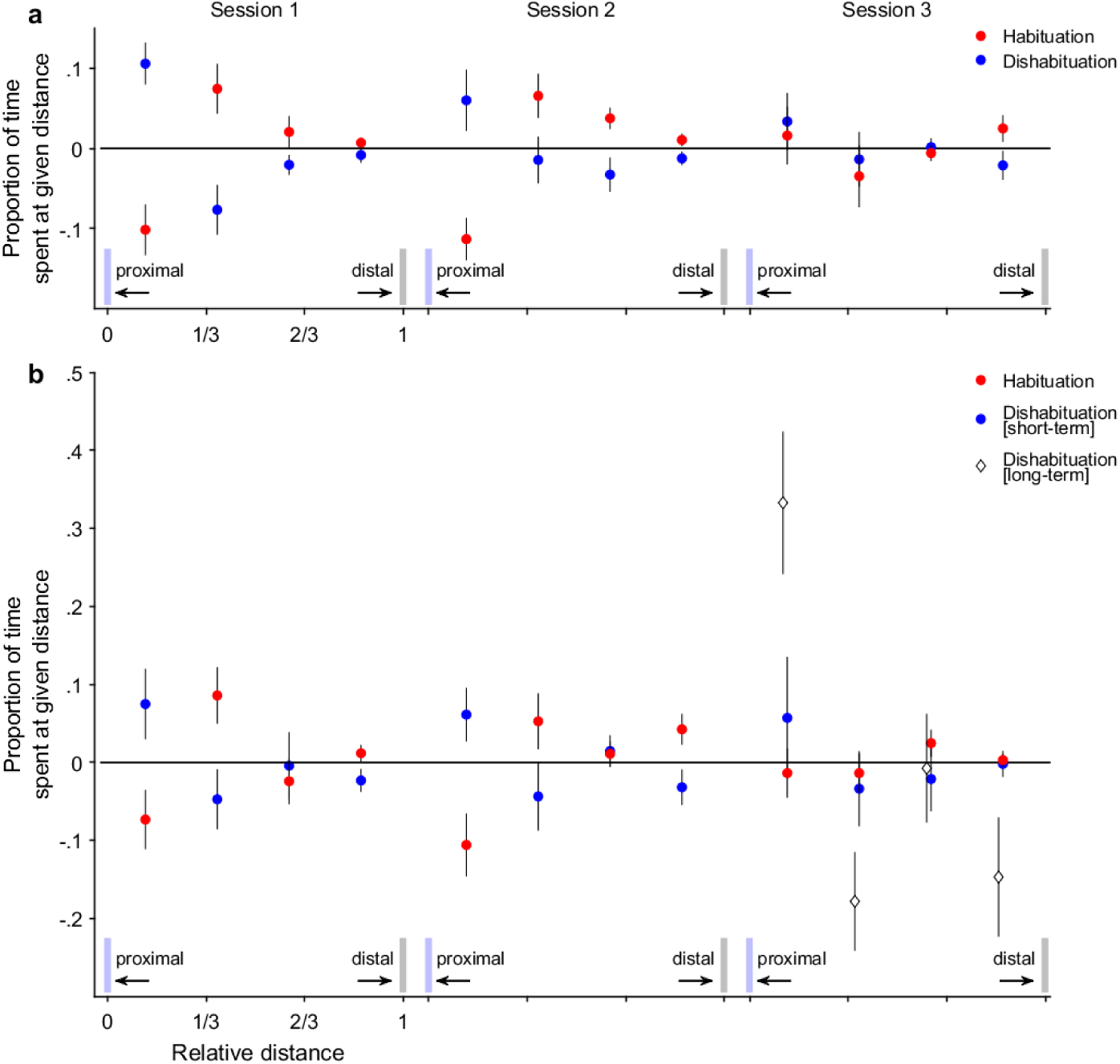
The relative change in distance between pairs of individuals, upon being confronted with the same individual as in the preceding trial (habituation trial; red discs) or a different individual from the individual in the preceding trial (dishabituation trial; blue discs). Each panel refers to an experiment (panel a. for Experiment 1; panel b. for Experiment 2), consisting of three sessions of trials. The dependent data is shown as the proportion of time spent at a given distance binned into 4 equally spaced bins. The x-axis labels refer to the proportional distances from the transparent acrylic sheet, ranging from ‘proximal’ to ‘distal’; the y-axis refers to the proportion of time spent at a given distance, i.e. the relative number of samples that fall into a given bin. Discs show the mean proportion across all individuals (i.e. 20 for Experiment 1; 16 for Experiment 2). The whiskers indicate the standard errors of the mean. White diamonds in the lower right subfigure b show the long-term dishabituation trials. Light blue bars indicate the side of the transparent acrylic sheet (proximal); grey bars indicate the back wall of the container (distal)

Furthermore, the strength of this effect varied across sessions. The interaction between *distance* and *condition* was significantly modulated by *session*, indicating that the dissociative effect of *condition* changed over the course of the testing period. This effect was strongest in Session 1 and weakened progressively, with the weakest modulation observed in Session 3 (LRT: 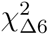 = 34.14, *p* < 0.001; Figure 2a, Supplementary Table 1, exemplar trial: Figure 1 d-f).

The systematic dissociation of distance values between *habituation* and *dishabituation* trials suggests that *P. regius* is capable of individual recognition. If spiders did not differentiate between conspecifics, we would not expect such a consistent divergence in distance patterns between conditions (*habituation* vs *dishabituation*). In addition, this dissociation diminished across sessions, indicating a progressive habituation effect towards the already encountered individuals. That is, in Session 1, the initial exposure to each individual elicited a pronounced dishabituation response. By Session 2, this dishabituation response was still present but attenuated. By Session 3, the dishabituation response had largely disappeared, indicating that spiders no longer differentiated between familiar and novel individuals (Video 1 and Supplementary Videos 1, 2 in the OSF repository (https://osf.io/gpnct/)). While this progressive reduction is consistent with the formation and retrieval of individual-specific memories, alternative explanations, such as general fatigue due to extended testing periods, cannot be ruled out at this stage.

As a second step, we therefore examined whether *P. regius*’s decreasing interest across repeated sessions is driven by a general fatigue effect due to the prolonged testing procedure, or whether *P. regius* recognises the current individual after having encountered it at least once (by Session 2) or twice (by Session 3), and as a result, no longer dishabituates. Such recognition capability would support the presence of long-term memory representations in the individuation of conspecifics, given the extended retention interval spanning from minutes to hours.

To this end, we assessed whether the presentation of a completely novel individual - unseen during any of the three experimental sessions - would elicit a rebound in interest when introduced at the end of Session 3. We refer to this condition as *dishabituation [long-term]* trials, in contrast to the *dishabituation* trials conducted during Sessions 1-3, henceforth referred to as *dishabituation [short-term]* trials (see Table 3). Importantly, the labels ‘short-term’ and ‘long-term’ do not imply specific memory systems, but rather reflect the retention interval at which the respective dishabituation trials were administered.

**Table 3:**
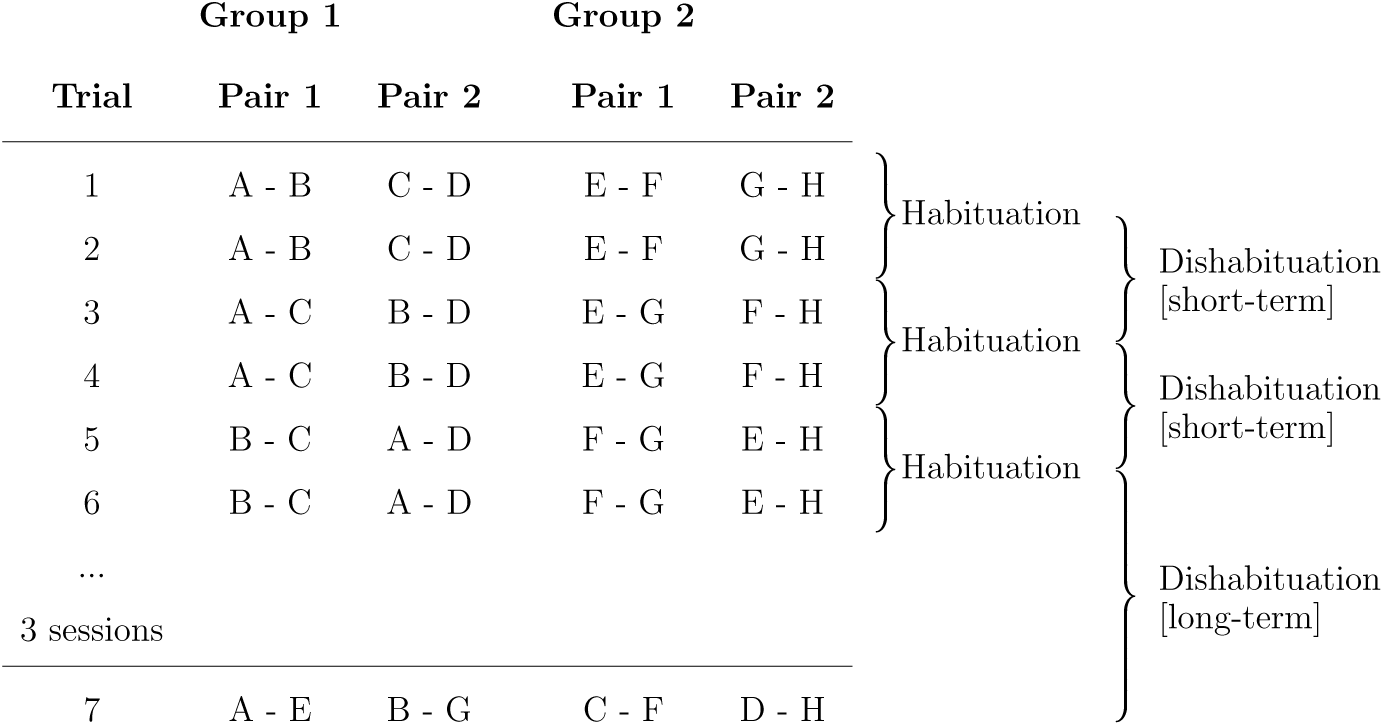
Pairwise comparisons as habituation and dishabituation trials.

If such rebound occurs, it would indicate that the observed habituation across sessions results from the recognition of repeatedly presented individuals, rather than from physical fatigue due to the prolonged testing procedure. In other words, such rebound would suggest a form of ‘cognitive’ fatigue - a diminished response elicited by the repeated re-encounter of familiar individuals - subserved by long-term memory formation, rather than a ‘physical’ fatigue effect. To test this, we re-ran the experiment in an additional 16 spiders, arranged to four groups and added a memory *dishabituation [long-term]* trial at the end of Session 3. The memory *dishabituation [long-term]* trials were generated by cross-combining individuals from two groups (group 1: A, B, C, D; group 2: E, F, G, H; Table 3) that had been run in parallel: At the end of Session 3, each spider was paired with a novel individual from the other group (e.g., A - E, B - G, C - F, D - H), resulting in previously unseen pairings.

First, we observed that spiders responded with renewed interest toward these unfamiliar individuals (*dishabituation [short-term]* trials), approaching them more closely than they had approached previously encountered ones (habituation trials). Thus, we replicated our earlier findings and found a dissociation between the factors *distance* and *condition* (LRT: 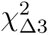 = 29.52, *p* < 0.001; Figure 2b, Video 2 and Supplementary videos 3, 4 in the OSF repository (https://osf.io/gpnct/), Supplementary Table 2). Specifically, *dishabituation [short-term]* trials (blue discs) showed a greater proportion of close-distance values, whereas *habituation trials* (red discs) showed a greater proportion of far-distance values. Most critically, we found that the *dishabituation [long-term]* trials at the end of Session 3 elicited a rebound in interest that clearly exceeded the rebound in the *dishabituation [short-term]* trials of the same session. This was reflected in a significant interaction between *condition* (i.e. *dishabituation [short-term]* vs *dishabituation [long-term]*) and *distance* (*F* (3,127) = 3.91, sum sq. = 0.92, mean sq. = 0.31, *p* < 0.01, Figure 2b (right subfigure, white diamonds; exemplar trials: Figure 1g-i, j-l, Videos 3 - 5 and Supplementary videos 5, 6 in the OSF repository (https://osf.io/gpnct/)). This finding suggests that the reduction in effect size across sessions was not due to general fatigue, but rather reflects a long-term memory for previously encountered individuals: That is, when presented with a truly novel conspecific, the spiders’ interest rebounded to an unprecedented level.

Our findings show, first, that *P. regius* can recognise individuals to which it was exposed to for a short period of 7 minutes and that reoccurred after a visual separation period of 3 minutes. Second, *P. regius* exhibited long-term habituation, i.e. a sustained reduction in interest when encountering the same individuals again 1 or 2 hours later. This result pattern is only possible if spiders remember the encountered individuals across sessions, suggesting the formation and retention of individual-specific memory representations. Third, despite this long-term habituation, *P. regius* showed an unprecedented rebound in interest when confronted with an entirely novel individual, ruling out a physical fatigue effect in favour of a ‘cognitive’ fatigue based on long-term memory. For these reasons, our results are the first evidence that *P. regius*, a non-social arthropod species, possesses long-term memory that allows it to individuate conspecifics and recognise novel individuals.

Before addressing the evolutionary implications of these findings, it is important to consider a key methodological question: Do the observed behavioural changes reflect true recognition memory, or could they result from simpler mechanisms such as transient familiarity or physical fatigue? To distinguish these possibilities, we employed a stringent habituation–dishabituation paradigm [15], widely used in animal [14] and infant cognition [16] to isolate memory-based recognition from short-term novelty effects. Unlike paradigms with static images or single exposures, our approach required subjects to recognise a dynamic, interacting conspecific across repeated encounters, making low-level novelty detection or general desensitisation unlikely explanations. In our design, evidence for individual recognition comes from both within-session and across-session comparisons: Within sessions, *dishabituation [short-term]* trials consistently elicited closer proximity-seeking behaviour than *habituation* trials, indicating discrimination between familiar and novel individuals. Across sessions, this effect showed a structured decline: strong in Session 1, attenuated in Session 2, and absent in Session 3. This temporal pattern is inconsistent with a general desensitisation, which would reduce proximity-seeking behaviour across all trial types regardless of stimulus identity. Instead, our results support the formation of identity-specific memory representations. Further support comes from the long-term dishabituation trials at the end of Session 3, where introducing a novel individual elicited the strongest rebound in interest. If prior responses reflected transient familiarity or physical fatigue, such a pronounced effect would not be expected. This indicates that spiders encoded previous individuals as distinct representations and recognised novelty at the level of individual identity.

Our results provide clear evidence of memory-based individual recognition in *P. regius*. If a spider does not dishabituate when re-encountering an individual, this indicates that a memory representation for that individual has been formed and retrieved. Conversely, if a marked dishabituation response occurs upon introduction of a novel individual, it implies that the current experience does not match any existing memory representation, prompting renewed interest. The clear dissociation between habituation and dishabituation trials within sessions, the progressive reduction in dishabituation responses across sessions, and the pronounced rebound upon introduction of a truly novel individual together indicate the formation and retrieval of individual-specific memories, rather than transient familiarity or physical fatigue.

Recognising members of one’s own species is a crucial cognitive ability that underpins various adaptive behaviours. Individual recognition allows animals to distinguish between friend and foe, to identify a mating partner, its offspring or a kin member. Individual recognition is achieved via the production of individually-distinct features (e.g. visual) or signals (e.g. acoustic) by the sender, and the ability of the receiver to extract and process these features and signals [17]. In social species, individual recognition bears particular significance in contexts such as territoriality, aggressive competition and parental care [11]. It is precisely because jumping spiders, including *P. regius*, are generally considered non-social, solitary, and aggressive towards conspecifics that the presence of individual recognition is unexpected. This raises the question about the biological relevance of individual recognition in *P. regius*: One of the few social interactions in the life of a jumping spider occurs during mating, encompassing a highly structured visual courtship display composed of coordinated body movements and distinct morphological features. It is believed that the colouration of the appendages (the chelicerae), along with the facial hair patterns serve as important visual features for species and sex identification in jumping spiders, and may also signal individual quality during mate assessment [18, 19]. Hence, the colouration of body parts (sender) and the ability to discriminate colours (receiver) appear sufficient for sexual selection, such as species and sex identification or mate assessment, without necessarily supporting individual recognition [20, 18]. Similarly, in aggressive interactions, such as territorial disputes, fighting ability is largely associated with the size and colouration of the chelicerae [20, 21]. In line with this, research has shown that the demands of mating and aggression in jumping spiders are met by cue recognition - by definition a basic-level classification - allowing classification of species, sex, or general rival status. For example, jumping spiders have been shown to use a combination of visual and chemical (pheromonal) cues to differentiate both species and sex, enabling them to identify potential mates or competitors [9]. Some species recognise rivals by relying on distinctive colour signals, such as a red facial patch, which serves as a key visual cue in aggressive interactions [10]. In each case, it is basic-level category recognition rather than individual identity that guides behaviour, and thus subordinate-level discrimination appears unnecessary for these core ecological functions. As such, territoriality and aggressive behaviour appear insufficient as ultimate explanations [22] for the evolution of individual recognition in *P. regius*. Instead, decisions in these contexts can be based on general physical features rather than identity-specific information. In addition, most jumping spiders exhibited limited parental care [23], protecting the nest through the spiderlings’ first molt. A notable exception is *Toxeus magnus*, which provides a nutrient-rich, milk-like substance to its offpsring, a behaviour functionally and behaviourally analogous to lactation in mammals [24]. Despite these examples, there is little evidence that memory-based recognition of individual offspring is required to support parental care in salticids. In more specific contexts, however, some jumping spiders have been shown to discriminate their own eggsacs from those of conspecifics, presumably to avoid filial cannibalism [25], or to distinguish their own silk draglines from those of other conspecifics, representing a form of self-recognition [26]. Both forms of recognition, however, are context-dependent and do not extend to memory-based identification of conspecifics across repeated encounters. Hence, although jumping spiders demonstrate a range of basic-level recognition abilities adapted to specific ecological challenges, these abilities, however, do not extent to the subordinate-level individual recognition demonstrated in the present study. In other words, even in this context, there is no clear evidence that memory-based individual recognition solves a critical survival problem.

Nevertheless, it is worth considering that individual recognition could provide adaptive benefits in certain social contexts, even in taxa where repeated encounters are typically rare. For instance, the so-called ‘dear enemy effect’, a phenomenon observed in territorial animals, involves reduced aggression towards familiar neighbours while maintaining heightened vigilance against unfamiliar individuals [27, 28]. This form of recognition can minimise the costs of unnecessary conflict and enhance social stability. While the natural history of *P. regius* suggests rather infrequent repeated encounters with the same conspecific, the current findings raise the intriguing possibility that cognitive capacities for identity recognition facilitate the emergence of ‘dear enemy’ relationships if territorial dynamics arise. Future experiments could directly test whether the ‘dear enemy effect’ can be experimentally elicited in jumping spiders.

Taken together, the generally solitary nature of *P. regius* offer little ecological pressure to favour individual recognition. According to the ‘minimum needs hypothesis’ [29, 2], selection will only favour specific recognition abilities when basic-level classification [30] (e.g. species, sex, or quality assessment via colouration or size cues) no longer suffices to solve ecologically relevant problems. In the case of *P. regius*, basic discrimination, such as chelicerae size or colouration, may be sufficient in most known contexts of social interaction. Moreover, the neural implementation of subordinate-level classification [30], such as identity recognition, requires more detailed and exhaustive processing than basic-level categorisation. This type of fine-grained discrimination is typically associated with specialised neural structures and increased cognitive and neural resource requirements [31]. Given these costs and the limited ecological need for identity recognition in *P. regius*, it is more plausible to assume that this ability arises not through direct selection, but through pleiotropy, as a byproduct of general cognitive architecture evolved for other purposes - a view consistent with the ‘general learning hypothesis’ [29]. Jumping spiders are known for complex foraging and navigation strategies [32, 33, 34], requiring flexible learning and high behavioural adaptability. These domain-general cognitive capabilities may enable recognition of conspecifics with a degree of detail and specificity that exceeds their minimum recognition needs [17].

In contrast to *social* animal species, including eusocial arthropod species [6, 4, 7], where individual recognition is often maintained by direct selective advantage, we cannot conclusively identify a survival benefit for individual recognition in *P. regius*. Instead we put forward the idea that individual recognition in *P. regius* emerges as a pleiotropic consequence, namely a byproduct of an already fairly sophisticated, broader learning capability. Critically, individual recognition relies on recognition memory, a form of long-term memory, in which a previously encountered event or entity, here an individual, is stored as a representation and reactivated upon re-encounter [35]. This memory mechanism may also underlie complex behaviours observed in other salticids, such as when *Portia fimbriata* navigates towards prey after an initial visual scan of the environment - even without visual input during execution [36] - or when *Menemerus semilimbatus* recognises biological motion, demonstrating sophisticated temporal integration and dynamic cue recognition [37].

Thus, our study challenges the notion of spiders being stimulus-response driven automata, by not only contributing to an increasing body of evidence that spiders and saliticids in particular produce a wide spectrum of intelligent behaviour [38], but by pinpointing the presence of two fundamentally important mechanisms for any higher cognitive processing: flexible learning and recognition memory. The key building blocks of these mechanisms are representations, mental images of external entities, that are not present to the sense organs, allowing more elaborate information processing, such as in complex decision making and goal-directed behaviour. The existence of which in arthropods in general and spiders in particular triggers rethinking of miniature brain cognition [38].

## Materials and methods

### Subjects

Our subjects were 36 jumping spiders (*Phidippus regius*), kept individually in enclosures (7 x 7 x 12 cm) at room temperature (21 - 25°C) and supplied with a moist water-pad, exchanged every other day, and two small-sized cockroaches (*Shelfordella lateralis*) per week. All spiders were adult laboratory-bred and had no direct encounters with conspecifics during adulthood. Behavioural enrichment [39] was provided by means of climbing and nesting structures (i.e., natural wood branch) and by interaction with human caretakers and experimenters during handling and maintenance procedures. In Experiment 1, spiders were assigned to five experimental groups, three of which contained females, two of which males; in Experiment 2, spiders were assigned to four experimental groups, i.e. two groups per sex.

### Apparatus

In the following we describe how pairs of spiders were brought into direct visual contact under controlled conditions and in a manner that allows to reassign individuals easily and without interruption to form novel pairings. To this end, we built a cubical experimental arena of 60 by 46 by 65cm (L x W x H), consisting of white polypropylene plastic panels, mounted in a frame of T-slotted aluminium profiles (20 Series; Misumi Group Inc., Bunkyo City, Tokyo, Japan). Two LED light sources (Mettle^®^ SL400, 45W, 2100lm, 350 x 250mm surface area, Mettle Photographic Equipment Corporation, Changzhou, China) were placed outside the cubicle at 25cm distance from the side panels of the cubicle, illuminating the inside of the cubicle uniformly. We also mounted two FLIR^®^ 1.3MP, Mono Blackfly USB3 cameras with a 1/2” CMOS sensor (BFS-U3-13Y3M-C, FLIR^®^ Integrated Imaging Solutions, Inc, 12051 Riverside Way, Richmond, BC, Canada) equipped with 8mm UC Series lenses from Edmund Optics^®^ (Stock #33-307, Edmund Optics^®^, Barrington, New Jersey, USA) on T-slotted aluminium profiles, facing downwards onto the arena surface at a distance of 60cm. For each spider we 3D-printed a white container with outer dimensions (L x W x H) of 7 x 7 x 5cm and inner dimensions of 6.3 x 6.3 x 4.5cm. The upper side of the container and one of the four side walls were made of a transparent .5mm thick acrylic sheet. While the acrylic sheet on the upper side of the container was screwed onto the side walls of the container, the acrylic sheet on one of the sides of the container can be lifted up to open the container, allowing easier transfer of the spider from the home enclosure.

### Procedure

In Experiment 1, each group consisted of four same-sex spiders, with each spider being placed inside a container prior to the experiment. We allowed the spiders sufficient time (10 - 15min) to acclimatise to the new environment. During the experiment, the spiders remained in their own containers. With start of the experiment, we then placed the containers of the four spiders such that the transparent side walls of two containers were facing each other, resulting in two pairs of spiders with direct visual contact to each other. During the arrangement of the containers and before each new trial, visual contact was blocked using an occluder placed between the transparent side walls of the containers.

Each trial was initiated by removing this occluder, allowing visual contact. For simplicity, let the four individuals be symbolised by the letters ‘A’, ‘B’, ‘C’, and ‘D’: An arrangement of trials where each individual is opposed to each other individual is described in Table 1.

To tease apart, whether or not *P. regius* was capable of visually discriminating other individuals two types of trials were required: (a) a *habituation* trial, where the same individual was presented in the trial preceding the current trial (e.g., trial 1: A - B, trial 2: A - B), and (b) a *dishabituation* trial, where a different individual was presented in the trial preceding the current trial (e.g., trial 1: A - B, trial 2: A - C). Thus, each *dishabituation* trial followed a *habituation* trial, forming alternating *habituation* and *dishabituation* phases, respectively, as shown in Table 2.

In detail, a trial, e.g. A - B (and in parallel C - D), lasted for 7 minutes allowing the spiders to visually inspect each other, before isolating the spiders visually for 3 minutes with a non-transparent white occluder, fully covering the transparent side wall. Following the occluder phase, another 7-minute exposure phase was initiated, which consisted of either the same individual (*habituation* trial) or another individual (*dishabituation* trial) than in the preceding trial. During each trial, the distance between individuals was measured at 10Hz temporal resolution, and used as and indicator of ‘interest’: shorter distances signalled greater interest in the other individual, while longer distances indicated reduced interest.

This experimental design was specifically structured to assess identity recognition memory in *P. regius*. With the repetition of trials, *habituation* and *dishabituation* phases, we can assess whether a currently visually encountered individual (*habituation*) elicits a differential behavioural response to a novel individual (*dishabituation*). We predict a dissociation of distances between *habituation* and *dishabituation* trials. This habituation - dishabituation paradigm is widely used in developmental [16] and animal cognition studies [14] to evaluate recognition memory in non-verbal subjects.

With the outlined procedure (Table 2), we can form sequences of trial-phases, where each first of two trials is a habituation trial, and every second is a dishabituation trial. Thus, a *habitation* trial consists of a repeated pairing, such as A - B followed by A - B, while a *dishabituation* trial consists of a novel pairing, such as A - B followed by A - C. As a result, each trial within a trial-phase subserves both the *habituation* as well as the *dishabituation* trial. In this manner, we created a trial list, containing 12 trials in total, six of which result in *habituation* trials and six of which result in *dishabituation* trials (Table 2). This session of trials was repeated twice, resulting in a total of 36 trials per experiment. Each experiment lasted 180 min, where each trial contained 7 min exposure and 3 min visual separation. Each group of spiders was subjected to this protocol.

Two amendments were made in Experiment 2: (a) we ran two groups of four individuals in parallel, and (b) we introduced additional *cross-group* trials were introduced at the end of Session 3. This resulted in a modified procedure described in Table 3.

### Data logging and analysis

Camera control and image acquisition were done using Matlab (Mathworks^®^, Natick, Massachusetts, USA) and the image acquisition and processing toolboxes. The frame rate was set to 10Hz. Cameras were placed perpendicular to the xy-plane at a distance of about 60cm from the ground. The lens aperture was set to f/4, allowing a sufficient depth of field. Analysis was done with Matlab (Mathworks^®^, Natick, Massachusetts, USA). We pre-processed the video recordings by segmenting the spider body from the background in each frame using functions for image intensity adjustment, image enhancement, image binarization and image properties measurement to extract the largest available ‘region’, the spider body, and its centroid. For each trial we approximated the distance between the individuals in the xy-plane as a function of time, using the Euclidean distance weight function based on the centroid coordinates of the two individuals. We then pooled the distance values of each trial into 4 equally-sized and non-overlapping bins (bin centers [mm]: [20, 60, 100, 140]; bin size 40mm; maximal distance « 160mm) and calculated the proportion of time spent at a given distance. Each bin was normalised by the total number of events. Differences between proportions were then calculated for every trial comparison according to Tables 2, 3: For instance, the proportions of time spent at a given distance for individual A in trial 1 was subtracted from the proportions of time spent at a given distance for individual A in trial 2, resulting in an assessment for the relative rebound of interest following a repetition of exposure to the same spider B (*habituation*). Subsequently, the proportions of time spent at a given distance for individual A in trial 2 was subtracted from the proportions of time spent at a given distance for individual A in trial 3, resulting in an assessment for the relative rebound of interest following changes in spider’s identity (*dishabituation*). We used linear mixed-effects models, where the differences in proportions served as the dependent variable. We fitted two separate models for each experiment (*Full model 1* and *2*), and followed a commonly accepted model fitting procedure [40]: To fully account for the dependent variable, we fitted three predictor variables: (1) The bin number ([1 to 4]), reflecting a discretised distance measure and henceforth referred to as factor *distance* ([1 to 4]), (2) the *session* of comparisons ([1, 2, 3]), as outlined in the Procedure above (Table 2, 3), and (3) the *condition*, referring to whether the given comparison was a *habituation* or *dishabituation* comparison. We also fitted all two-way interactions between the three main predictors: *distance*:*session*, *distance*:*condition*, and *session*:*condition*, as well as the three-way interaction *distance*:*session*:*condition*. Of particular interest are the two-way interaction between the factors *distance* and *condition*, since we predict a modulation of *distance* values by *condition* as a function of *distance*, and the three-way interaction between the factors *distance*, *condition* and *session*, since we predict a modulation of *condition* as a function of *distance* which becomes weaker over time and repetitions, i.e. *session*. We further defined *sex* of the subjects and *subject* as random factors in all models. We fitted a linear mixed-effects model (fitlme function in Matlab) with normal error structure and identity link function to our data set. We then created a null model for each corresponding full model, which consisted of the similar structure as the full model, however leaving only *distance* as fixed effect, while preserving all random effects. Using likelihood ratio test (LRT), we compared the null models with the corresponding full models. Assuming a significant improvement for the full model over the null model, the non-significant interaction terms were removed from the full model, reaching a model containing only significant interaction terms and both significant and non-significant main effects [41, 42], henceforth referred to as the final model. Evaluation of fixed effect were on the basis of the final models and are referred to as the *Final model 1* (Experiment 1), and *Final model 2* (Experiment 2). This procedure resulted in the following models (Wilkinson notation):

*Final model 1 and 2 :*

’Response „ 1 + Distance + Session + Condition + Distance:Condition + … Distance:Session:Condition + (1|Sex) + (1|Subject)’

An additional analysis of variance was performed comparing the *dishabituation [long-term]* trials at the end of Session 3 with the *dishabituation [short-term]* trials from Session 3 (Table 3) as a function of *distance*. No statistical methods were used to predetermine sample size. The experiments were not randomized. The investigators were not blinded to allocation during experiments and outcome assessment.

## Acknowledgements

We are grateful for the financial support by the Swiss National Science Foundation (PZ00P3 154741), the Startup-funding of Taipei Medical University (TMU108-AE1-B33) and the Taiwan Ministry of Science and Technology research grants (110-2311-B-038-002, 112-2410-H-038-027) awarded to CDD. We thank Guillaume Dezecache, Niall W. Duncan, Olivier Pascalis, Tzu-Yu Hsu, Werner Müller and Timothy J. Lane for suggestions and comments on the manuscript.

## Author contributions

CDD: study design, data collection, analysis and interpretation, writing article, provision of necessary tools; YC: data collection, writing article, provision of resources.

## Additional information

All authors have seen and approved the manuscript. The manuscript has not been accepted or published elsewhere. Supplementary Information is available for this paper. Correspondence and requests for materials should be addressed to Christoph D. Dahl. Codes and materials are available (https://osf.io/gpnct/).

## Ethical approval

According to Taiwan’s Animal Protection Act, issued by the Council of Agriculture (Executive Yuan), experiments on invertebrates are allowed to be conducted without any special permission in Taiwan.

