## Supplementary information for "Individual recognition in a jumping spider (*Phidippus regius*)"

### Supplementary material

#### Materials and methods

Model parameter estimations for experiments 1 and 2, as described in the maintext, can be found in the Tables 1 and 2 below. An evaluation of the residuals of the final models can also be found below (Supplementary Figure 1). Supplementary videos are available, showing individual trial combinations of *habituation* and *dishabituation* trials. This typically includes a *baseline* trial (shown in the left panel of subfigure a), which is the *dishabituation* trial of the previous comparison. Also, see Table 2 in the article for clarification of the experimental design.

### Tables

*Table 1: Results of the model investigating pairwise subtracted frequency distribution of distance values. The table contains parameter estimates for the final model 1 based on the fixed factors 'distance', 'session', 'condition', 'distance:condition', 'distance:session:condition', as well as the random factors 'sex' and 'subject'.*

| | | **Estimate** | **SE** | ***t*-stat** | **DF** | ***p*-value** | **CI (95%) [lower, upper]** |
|---|---|---|---|---|---|---|---|
| | Intercept | -0.001 | 0.01 | -0.001 | 1424 | 1 | [-0.01; 0.01] |
| **Distance** | Distance 1 | 0.001 | 0.01 | 0.001 | 1424 | 1 | [-0.02; 0.02] |
| | Distance 2 | 0.001 | 0.01 | 0.001 | 1424 | 1 | [-0.02; 0.02] |
| | Distance 3 | -0.001 | 0.01 | -0.001 | 1424 | 1 | [-0.02; 0.02] |
| | (against *Distance* 4) | | | | | | |
| **Session** | Session 1 | 0.001 | 0.01 | 0.001 | 1424 | 1 | [-0.01; 0.01] |
| | Session 2 | 0.001 | 0.01 | 0.001 | 1424 | 1 | [-0.01; 0.01] |
| | (against *Session* 3) | | | | | | |
| **Con-dition** | Condition 1 | 0.001 | 0.01 | 0.001 | 1424 | 1 | [-0.01; 0.01] |
| | (against *Condition* 2) | | | | | | |
| **Distance x Condition** | Distance 1 : Condition 1 | -0.07 | 0.01 | -7.90 | 1424 | 0.001 | [-0.08; -0.05] |
| | Distance 2 : Condition 1 | 0.03 | 0.01 | 4.16 | 1424 | 0.001 | [0.02; 0.05] |
| | Distance 3 : Condition 1 | 0.02 | 0.01 | 2.06 | 1424 | 0.05 | [0.01; 0.03] |
| | (against *Distance* 4 and *Condition* 2) | | | | | | |
| **Distance x Session x Condition** | Distance 1 : Session 1: Condition 1 | -0.04 | 0.01 | -3.14 | 1424 | 0.01 | [-0.06; -0.01] |
| | Distance 2 : Session 1: Condition 1 | 0.04 | 0.01 | 3.41 | 1424 | 0.001 | [0.02; 0.06] |
| | Distance 3 : Session 1: Condition 1 | 0.01 | 0.01 | 0.27 | 1424 | 0.78 | [-0.02; 0.02] |
| | Distance 1 : Session 2: Condition 1 | -0.02 | 0.01 | -1.71 | 1424 | 0.08 | [-0.04; 0.01] |
| | Distance 2 : Session 2: Condition 1 | 0.01 | 0.01 | 0.42 | 1424 | 0.67 | [-0.02; 0.03] |
| | Distance 3 : Session 2: Condition 1 | 0.02 | 0.01 | 1.50 | 1424 | 0.13 | [-0.01; 0.04] |
| | (against *Distance* 4, *Session* 3 and *Condition* 2) | | | | | | |

*Table 2: Results of the model investigating pairwise subtracted frequency distribution of distance values. The table contains parameter estimates for the final model 2 based on the fixed factors 'distance', 'session', 'condition', 'distance:condition', 'distance:session:condition', as well as the random factors 'sex' and 'subject'.*

*Table 2: Results of the model investigating pairwise subtracted frequency distribution of distance values. The table contains parameter estimates for the final model 2 based on the fixed factors 'distance', 'session', 'condition', 'distance:condition', 'distance:session:condition', as well as the random factors 'sex' and 'subject'.*

| | | **Estimate** | **SE** | ***t*-stat** | **DF** | ***p*-value** | **CI (95%) [lower, upper]** |
|---|---|---|---|---|---|---|---|
| | Intercept | 0.01 | 0.01 | 0.01 | 1136 | 1 | [-0.01; 0.01] |
| **Distance** | Distance 1 | -0.01 | 0.01 | -0.01 | 1136 | 1 | [-0.02; 0.02] |
| | Distance 2 | 0.01 | 0.01 | 0.01 | 1136 | 1 | [-0.02; 0.02] |
| | Distance 3 | 0.01 | 0.01 | 0.01 | 1136 | 1 | [-0.02; 0.02] |
| | (against *Distance* 4) | | | | | | |
| **Session** | Session 1 | 0.01 | 0.01 | 0.01 | 1136 | 1 | [-0.02; 0.02] |
| | Session 2 | -0.01 | 0.01 | -0.01 | 1136 | 1 | [-0.02; 0.02] |
| | (against *Session* 3) | | | | | | |
| **Con-dition** | Condition 1 | -0.01 | 0.01 | -0.01 | 1136 | 1 | [-0.01; 0.01] |
| | (against *Condition* 2) | | | | | | |
| **Distance x Condition** | Distance 1 : Condition 1 | -0.06 | 0.01 | -5.14 | 1136 | 0.001 | [-0.09; -0.04] |
| | Distance 2 : Condition 1 | 0.04 | 0.01 | 3.32 | 1136 | 0.001 | [0.02; 0.07] |
| | Distance 3 : Condition 1 | 0.01 | 0.01 | 0.30 | 1136 | 0.77 | [-0.02; 0.03] |
| | (against *Distance* 4 and *Condition* 2) | | | | | | |
| **Distance x Session x Condition** | Distance 1 : Session 1: Condition 1 | -0.01 | 0.02 | -0.55 | 1136 | 0.58 | [-0.04; 0.02] |
| | Distance 2 : Session 1: Condition 1 | 0.02 | 0.02 | 1.41 | 1136 | 0.16 | [-0.01; 0.06] |
| | Distance 3 : Session 1: Condition 1 | -0.01 | 0.02 | -0.77 | 1136 | 0.44 | [-0.05; 0.02] |
| | Distance 1 : Session 2: Condition 1 | -0.02 | 0.02 | -1.09 | 1136 | 0.28 | [-0.05; 0.02] |
| | Distance 2 : Session 2: Condition 1 | 0.01 | 0.02 | 0.37 | 1136 | 0.71 | [-0.03; 0.04] |
| | Distance 3 : Session 2: Condition 1 | -0.01 | 0.02 | -0.31 | 1136 | 0.75 | [-0.04; 0.03] |
| | (against *Distance* 4, *Session* 3 and *Condition* 2) | | | | | | |

### Figures

Supplementary Figure 1

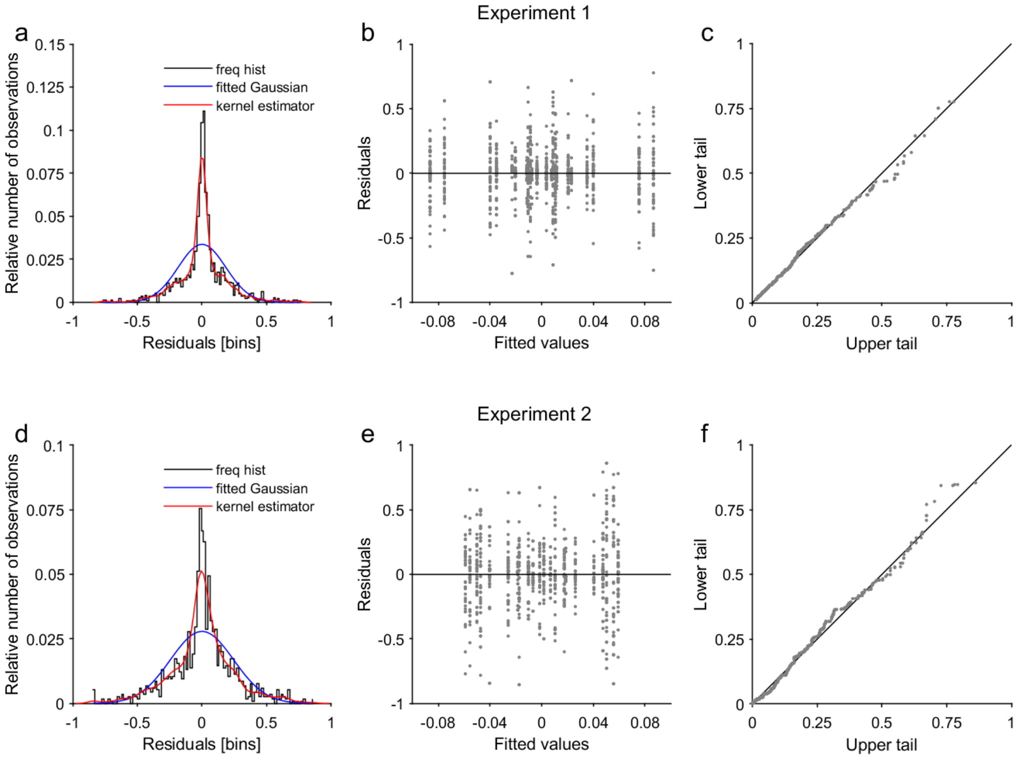


**Supplementary figure caption 1**: Residuals of the linear mixed-effects model. a, d. Residuals of linear mixed-effects model 1 (a) and 2 (d). The histograms bin the residuals for each model into 100 equally-spaced containers (x-axis) and return the number of elements (y-axis) in each container. The blue lines indicate a Gaussian fit; the red lines show a kernel distribution estimate. b, e. show the residuals (y-axis) plotted against the fitted values (x-axis) for model 1 (b) and model 2 (e). c, f. Residuals of lower and upper tails are plotted against each other, showing an equal distribution in both model 1 (c) and model 2 (f).

### Videos

*Supplementary videos*:

The videos can be found in the following repository: https://osf.io/gpnct/

The video speed is increased by a factor of approximately 5.6.

Dishabituation [short-term] trials **experiment 1**:

'supplementaryVideo01.avi'

'supplementaryVideo02.avi'

Dishabituation [short-term] trials **experiment 2**:

'supplementaryVideo03.avi'

'supplementaryVideo04.avi'

Dishabituation [long-term] trials:

'supplementaryVideo05.avi'

'supplementaryVideo06.avi'

**Supplementary video caption**: Example videos. a. Baseline, habituation and dishabituation trials are shown for a given individual (top half of the black box) with varying partner according to condition (lower half of the black box), i.e. baseline and habituation trials require an exposure to an identical partner; dishabituation trials to a different partner. Distances are indicated by a coloured dashed line (red for baseline, green for habituation, blue for dishabituation trials). The light blue lines dividing the black boxes illustrate the approximate placement of the transparent acrylic sheets. The black boxes indicate the approximate location of the walls of the containers. The exact location of walls and transparent front panels might slightly vary. The $\Delta$-values show the current relative distance between individuals for a given condition and $\bar{\mathbf{x}}$-values the mean relative distance up to the current sample. The maximal distance, i.e., when both spiders are in diagonally opposite corners, is 1, the minimal distance is 0. b. The distribution of relative distances of data samples up to the current point in time is shown as proportion of time spent at a given distance according to the trial types (baseline, habituation, dishabituation). The discs indicate the bin to which the current distance values are assigned to, and, hence, dynamically change their location as the spider moves. c. Trial types are shown as subtraction from each other, such that habituation trials are contrasted with baseline trials (black line), and dishabituation trials with habituation trials (dashed black line). Similarly to b., the discs indicate to which bin the current sample is assigned to. The vertical positioning of the discs indicate by their colours which trial type is more frequent at a given point, e.g., a blue disc located above a green disc indicates that for the given bin the dishabituation trial (blue disc) showed more values falling into that bin than the habituation trial (green disc) (≡ positive value); a green disc located above a blue disc indicates that for the given bin the habituation trial (green disc) showed more values falling into that bin than the dishabituation trial (green disc) (≡ negative value).
